# Photosynthetic capacity in seagrass seeds and early-stage seedlings of *Zostera marina* L

**DOI:** 10.1101/2022.12.12.520078

**Authors:** Kasper Elgetti Brodersen, Michael Kühl

## Abstract

**Summary:** In many terrestrial seeds, photosynthetic activity supplies O_2_ to the developing plant embryo to sustain aerobic metabolism and enhance biosynthetic activity. However, whether seagrass seeds possess similar photosynthetic capacity to alleviate intra-seed hypoxic stress conditions is unknown.

We used a novel combination of microscale variable chlorophyll fluorescence imaging, a custom-made O_2_ optode microrespirometry system, and planar optode O_2_ imaging, to determine the O_2_ microenvironment and photosynthetic activity in developing seeds and seedlings of seagrass (*Z. marina* L.).

Developing, sheath-covered seeds exhibited high O_2_ concentrations in the photosynthetic active seed sheath and low O_2_ concentrations in the center of the seed at the position of the embryo. In light, photosynthesis in the seed sheath increased O_2_ availability in central parts of the seed enabling enhanced respiratory energy generation for biosynthetic activity. Early-stage seedlings also displayed photosynthetic capacity in hypocotyl and cotyledonary tissues, which may be beneficial for seedling establishment.

Sheath O_2_ production is important for alleviating intra-seed hypoxic stress and can increase endosperm storage activity improving the conditions for successful seed maturation and germination.

## Introduction

Seagrass plants are marine angiosperms that form important coastal ecosystems playing a key role for coastal protection, biodiversity of marine organisms, and sequestration of nutrients and carbon (Madsen et al. 2001; Bertelli & Unsworth, 2014; Fourqurean et al. 2012). Yet, seagrass meadows are declining worldwide due to climate change and other anthropogenic threats (Orth et al. 2006; Waycott et al. 2009), which has stimulated increasing research on seagrass habitat restoration (Katwijk et al. 2016). Seed-based restoration is of major interest due to the potential for restoring larger areas and at the same time ensuring high genetic diversity in re-established seagrass meadows (Marion & Orth, 2012; Katwijk et al. 2016; Cumming et al. 2017; Unsworth et al. 2019). However, there are still large knowledge gaps in the understanding of fundamental mechanism and processes influencing seed development, germination and survival (York et al. 2017). Seagrass seed production is initiated in flowering shoots in the warming waters during spring (Silberhorn et al. 1983). During seed development, the seeds are either i) released from the plant sedimenting directly onto the sea floor, or ii) dispersed via detached decaying reproductive shoots or in seed-containing spathes on the sediment surface (Harwell & Orth, 2002; Marion & Orth, 2012; Hosokawa et al. 2015). Most seeds settle within or close to the meadow of origin (Harwell & Orth, 2002; Hosokawa et al. 2015). For *Z. marina* plants growing in temperate regions, seed germination occurs over the autumn/winter, whereafter the young seedlings commence rapid growth during the following springtime with the increased light availability (Marion & Orth, 2012).

Photosynthetic activity during seagrass seed germination and seedling development has been demonstrated for non-dormant seagrass species like *Posidonia oceanica* and *Thalassia testudinum* (Celdran & Marin, 2011 and 2013; Celdran, 2017). Seed photosynthesis enhances *Posidonia oceanica* seedling (leaf and root) growth (Celdran & Marin, 2013), where the autotropic production complements stored carbohydrate reserves until the seedling’s photosynthetic capacity is capable of supporting its own carbon demands (Celdran & Marin, 2013). The *Posidonia* seed photosynthesis also seems to support the seedlings O_2_ demand, a mechanism that could be important for avoiding hypoxia during seed embryogenesis (Celdran et al. 2015). Low internal O_2_ conditions can limit respiration rates and seed photosynthesis could potentially alleviate this hypoxic stress through an O_2_ supported supply of respiratory energy (Celdran et al. 2015). *Posidonia* seed photosynthesis could thus help alleviate the O_2_ demand required to metabolize the carbohydrate reserves stored in the seeds (Celdran et al. 2015). However, a similar mechanism remains to be demonstrated in dormant seagrass seeds such as found in the key model seagrass species *Z. marina* L. These seeds differ substantially from *Posidonia* seeds both in terms of their architecture and life cycle (i.e., seed sheath, hard seed coat and dormancy of up to 12 months; Orth et al. 2000) and may, therefore, have higher similarity to seed development of terrestrial monocot plants.

Seeds of *Z. marina* L. are known to have distinct dormancy and a hard seed coat (Orth et al. 2000), but its functional roles in terms of seed protection, photosynthetic activity and overall architecture remain largely unknown. In terrestrial seeds (e.g., seeds of soybean and grains of barley and maize; Borisjuk et al. 2005; Rolletschek et al. 2004, 2005), the function of the seed coat has mainly been related to protective properties. However, the seed coat also functions as a connection to the external microenvironment by transferring environmental cues into the seed, which leads to adjustments of the metabolic activity in terrestrial seeds (Radchuk & Borisjuk, 2014). The epidermal cuticle of the seed coat in terrestrial plant seeds forms a physical barrier but does not impede the exchange of gases such as O_2_ and CO_2_ (Radchuk & Borisjuk, 2014).

Developing seeds of terrestrial plants have high respiratory needs, generating hypoxic to near anoxic internal conditions, which thus requires additional O_2_ supplies to sustain aerobic respiration (e.g., Rolletschek et al. 2004, 2005). Low internal O_2_ conditions may be beneficial to the seed during development as it can stimulate high carbon use efficiencies through increased bioenergetic efficiency of mitochondria (Radchuk & Borisjuk, 2014 and references therein), and hypoxic conditions could also play a major role in enabling the longevity of seeds and provide means for regulating local metabolic activity (Borisjuk and Rolletschek, 2009). However, nutrient transport and endosperm storage activity largely depend on sufficient energy supply from respiration, which is dependent on sufficient O_2_ availability that can be enhanced by photosynthesis in the surrounding chlorenchymatic layer (Radchuk & Borisjuk, 2014). Photosynthesis in the seed coat has been demonstrated in many terrestrial seeds, driving seed carbon fixation and O_2_ production (Rolletschek et al. 2004), often at much lower light conditions than in leaves (Radchuk & Borisjuk, 2014). Seed coat photosynthesis can thus relieve hypoxic stress and enhance the biosynthetic activity of the developing seed (Rolletschek et al. 2003, 2005). The main input of nutrients to the developing seed is, however, supplied by the mother plant leaf, where the seeds’ ability to sense light enable metabolic adaptation to often uneven nutrient flows (Radchuk & Borisjuk, 2014). Gradients of seed photosynthesis in terrestrial monocotyledonous plant species (e.g., barley) can lead to internal O_2_ concentration gradients from high concentrations of O_2_ in the photosynthetic active chlorophyll-containing pericarp (chlorenchymatic regions) to low O_2_ concentrations in the starchy endosperm (storage organ) and embryo (Tschiersch et al. 2011): the endosperm in monocots is thus specialized for storage activity.

Internal O_2_ deficiency in the developing seed can limit respiration, which can be alleviated by seed photosynthesis improving respiration and likely higher synthesis of storage compounds positively affecting the overall yield of the developing seed (Tschiersch et al. 2011; Galili et al. 2014). Storage products such as carbohydrates and proteins are vital during the initial stages of seed germination. High respiration rates in the seed coat along with low tissue permeability can lead to high internal CO_2_ concentrations that promote internal CO_2_ re-fixation and thus enable an improved seed carbon budget (Wager, 1974; Flinn, 1985: Araus et al. 1993; Vigeolas et al. 2003; Rolletschek et al. 2004). Rubisco activity during seed photosynthesis mediates CO_2_ re-fixation, but the contribution of seed photosynthesis to biomass production via net CO_2_ fixation is considered low (Goffman et al. 2004; Ruuska et al. 2004; Radchuk & Borisjuk, 2014). In *Z. marina* seeds, the seed sheath is green during the early-stage development (e.g., Infantes & Moksnes, 2018) indicating the presence of a chlorenchymatic layer with active photosynthesis surrounding the non-green endosperm (i.e., the assimilating storage organ) in immature *Z. marina* seeds. However, such photosynthetic capacity of the *Z. marina* seed sheath (i.e., similar to the pericarp) and its implications for internal O_2_ concentration gradients and availability in the early-stages of seed development, have not been investigated.

In this study, we used high-resolution variable chlorophyll fluorescence imaging, custom-made micro-chambers for photosynthesis and respiration measurements (employing O_2_ sensitive sensor foils), and chemical imaging via planar O_2_ optodes to investigate the photosynthetic capacity and internal O_2_ concentration gradients of *Z. marina* (seeds with and without an intact seed sheath) as a function of increasing irradiance. We compared such detailed metabolic activity measurements on seeds to similar measurements on early-stage seedlings. We hypothesized that immature seeds with sheath and early-stage seedlings exhibit photosynthetic capacity that can support growth of the developing seed/plant via an O_2_-induced supply of respiratory energy that positively affects biosynthetic activity. Mature seeds, on the other hand, likely have low internal O_2_ conditions (and thus low photosynthetic capability) to support the longevity of the seed.

## Materials and Methods

### Seagrass Seed Sampling and Storage

Flowering shoots of *Zostera marina* L. with seeds were collected at Julebæk, Helsingør, Denmark (see Brodersen et al., 2020 for detailed description of the sampling site). After harvesting, the flowering shoots were stored in a large continuously aerated and illuminated (14h light/10h dark cycle) aquarium with similar light, temperature and salinity conditions as at the sampling site (temperature = 20°C; salinity = 18; incident photon irradiance = 200 μmol photons m^-2^ s^-1^). Immature sheath-covered seeds were collected directly from the spathes. Mature seeds were collected from the bottom of smaller tanks after naturally being released from the spathes. Only the denser seeds, i.e., rapidly sinking to the bottom after physical resuspension, were used for this study. Seed germination (*i*.*e*., split of seed coat and hypocotyl extension) was achieved via a low salinity (drop from 30 to 10) and high temperature (increase from ∼ 9 to 13 °C) pulse (e.g., Xu et al., 2016; Yue et al., 2019), where after the early-stage seedlings (less than 5 cm; Marion & Orth, 2012) and harvested seeds were used for variable chlorophyll fluorescence imaging and O_2_ dynamics measurements (see below).

### Experimental Setup and Treatments

All seagrass seeds and early-stage seedlings were kept in filter-sterilized seawater (0.2 μm; temperature of 21°C, salinity of 18) during measurements. For measurements of photosynthetic activity: photon irradiances of 0, 9, 99, 137, 232 and 323 μmol photons m^-2^ s^-1^ (where the highest light intensity was only used for the sheath-covered seeds and the early-stage seedlings) was evenly provided by a fiber-optic tungsten halogen lamp equipped with a collimating lens (KL-2500LCD, Schott GmbH, Germany). The incident photon irradiance (PAR, 400-700 nm) was measured for different lamp light settings with a scalar irradiance mini-sensor (US-SQS/L, Walz GmbH, Germany) interfaced to a pre-calibrated photon irradiance meter (ULM-500, Walz GmbH, Germany). To exclude the potential contribution of photosynthetic epiphytes on the O_2_ production and consumption measurements, selected seeds (n=4) were surface-sterilized via submersion into a saline ∼ 1.05% hypochlorite solution for 30s, followed by 3 × 1 min rinses in filter-sterilized (0.2 μm) seawater prior to measurements (Blaabjerg and Finster, 1998; Brodersen et al., 2018). To mimic the natural *in situ* environment of the mature seeds and early-stage seedlings, all O_2_ dynamic measurements were done at low O_2_ conditions of < 30 μmol L^-1^ (e.g., Brodersen et al., 2017 & 2019; Schrameyer et al., 2018; Trevathan-Tackett et al., 2020).

### Variable Chlorophyll Fluorescence imaging

Variable chlorophyll fluorescence imaging was used to determine the photosynthetic capacity and activity of *Z. marina* L. seed sheaths, seeds with and without epiphytes, and seedlings. A pulse– amplitude–modulation (PAM)-based variable chlorophyll fluorescence imaging system (MINI-IPAM; Walz GmbH) was used with a blue LED for measuring and actinic light (Ralph et al. 2005). Before measurements, the downwelling photon irradiance levels of the actinic light for specific program settings were calibrated with a quantum meter equipped with a cosine sensor. Images of the maximum photochemical quantum yield of PSII (i.e., F_v_/F_m_) was calculated as (Schreiber, 2004): F_v_/F_m_ = (F_m_-F_0_)/F_m_, where Fm is the maximal fluorescence yield measured during a strong saturation pulse fully closing all PSII reaction centres and F_0_ is the minimal fluorescence yield prior to the saturation pulse (measured after a dark-adaptation period of ∼15 min). Average F_v_/F_m_ values were calculated by integrating over the seed’s sheath or small photosynthetic active sites on mature seeds themselves, separately (i.e., within defined regions of interest), followed by calculating an average of all the investigated seeds for the respective development stages. Rapid light curves (RLCs) of the seeds and seedling photosynthetic activity were measured using series of images of effective PSII photochemical quantum yields (YII) and relative photosynthetic electron transport rates (rETR) at increasing irradiance with 20 s incubation at each irradiance step (Ralph and Gademann, 2005; Trampe et al., 2011). The effective photochemical quantum yield of PS(II) was calculated as: Y(II) = (F_m_lll−F)/F_m_ (Genty et al., 1989) and provides a measure of the PSII photosynthetic capacity. The rETR was calculated from Y(II) and the actinic photon irradiance, E_d_ as rETR = Y(II) × E_d_ and represents a relative measure of the PSII electron transport rate (e.g., Beer et al., 1998), assuming a constant PSII absorption cross section during measurements. All rETR, F_v_/F_m_ and Y(II) yields were calculated using the system software (ImagingWin, Heinz Walz GmbH, Germany).

### Multicolour microscope variable chlorophyll fluorescence imaging

Variable chlorophyll fluorescence of seagrass seeds and seedlings was also investigated with a red-green-blue (RGB) pulse-amplitude-modulated (PAM) chlorophyll fluorescence imaging system (RGB Microscopy-IPAM, Heinz Walz GmbH, Germany) mounted on an epi-fluorescence microscope (model Axiostar plus FL, Carl Zeiss, Jena, Germany; Trampe et al. 2011, Goessling et a. 2016) equipped with a 10x objective (Plan-Apochromate, Carl Zeiss GmbH, Germany). This microscope imaging PAM system allowed for high-spatial resolution measurements and analysis of single-cells and cross tissue distributions of photosynthetic capability. Relative photosynthetic electron transport rates (rETR), maximum (F_v_/F_m_) and effective quantum yields of PSII (YII) was determined with a white measuring light. The RGB fit function was used to investigate the composition of the seagrass seed epiphytic microbial community. In brief, the measuring principle of the RGB-fit is based on changes in fluorescence excitation spectra of different microbial oxyphototrophs, obtained by using sequential multicolour fluorescence imaging (i.e., red (R), green (G) and blue (B) excitation) and subsequent fluorescence image deconvolution into four different pigmentation types (Trampe et al. 2011). The analysis of the fluorescence excitation spectra differences between the actual measurements and a reference spectral matrix subsequently produces a colour-coded deconvoluted image, which enables identification of the abundance and distribution of diatoms, green algae, red algae and cyanobacteria. This is achieved as diatoms and green algae can be characterized by strong fluorescence excitation by blue light (460 nm) that overlaps with the absorption bands of Chl *b* and Chl *c*. Diatoms can thereafter be further distinguished by excitation with green light (525 nm) owing to presence of antenna such as fucoxanthin. Most cyanobacteria express maximal fluorescence yield with red–orange excitation (∼620 nm), owing to phycocyanin absorption, whereas red algae due to the presence of phycoerythrin display extraordinarily high fluorescence upon excitation with green light.

### Gas Exchange Measurements

Seed O_2_ production and consumption rates were measured in custom-made gas exchange chambers (1.8 mL) with pre-sterilized (0.2 μm) seawater employing calibrated O_2_ sensitive sensor spots (OXSP5, PyroScience GmbH, Aachen, Germany) connected to a fiber-optic O_2_ sensor system (FireSting O2, PyroScience, Germany), which was interfaced to a PC running dedicated data acquisition software (O_2_ logger, PyroScience, Germany). The oxygen sensor spots were 2-point calibrated in air saturated (100%) and anoxic (0%; obtained by flushing with N_2_ and by using an O_2_ scavenger such as sodium sulfite) seawater at experimental temperature and salinity. Water movement in the micro-photosynthesis and respiration chambers was maintained with glass coated magnets (www.sigmaaldrich.com: Spinbar® Pyrex® Glass-coated magnetic stirring bars) controlled by a magnet stirrer (IKA® big-squid; VWR International, Pennsylvania, USA). Glass coated magnets were used to avoid potential O_2_ consumption and/or release from the magnets during measurements. Closed gas exchange chambers (i.e., control and seagrass seed/seedling samples) were submerged into a small seawater tank, wherein the O_2_ concentration could be manipulated via flushing with N_2_. This setup constellation enabled replacing seeds and/or young seedling samples for biological replication without O_2_ intruding into the chambers when taking off the lid to change the samples. The O_2_ level in the small tank was monitored with a calibrated protected tip O_2_ minisensor (OXF500PT-OI; PyroScience, Germany) connected to a handheld fiber-optic O_2_ meter (FireSting GO_2_, PyroScience GmbH, Germany). Light at defined photon irradiance levels was provided by a fiber-optic tungsten halogen lamp equipped with a collimating lens (KL-2500LCD, Schott GmbH, Germany). We calculated gross photosynthesis, net photosynthesis and respiration rates from the measured O_2_ consumption or production rates as a function of incident irradiance as nmol O_2_ sample^-1^ h^-1^ (where controls served as blanks). In brief, the photosynthesis and respiration rates were calculated as follows: GP (E) = NP (E) + ⍰R(E)⍰, where respiration (R) and net photosynthesis (NP) rates were calculated from the linear O_2_ concentration slopes over time at the given photon irradiance (E), by multiplying the linear O_2_ concentration slope (nmol L^−1^ h^−1^) with the volume of seawater in the measuring chamber (L) divided by the sample wet weight. The gross photosynthesis (GP) rates were calculated by adding the absolute values of the respective respiration rates (i.e., dark respiration or post-illumination respiration; measured immediately after the light period at each irradiance level and used as a proxy for the respiration in the previous light period) to the respective net photosynthesis rates (further described in Hansen et al. 2022).

### Planar optode based O_2_ imaging

Planar O_2_ optodes with an isolating carbon black layer (Glud et al. 1996; Brodersen et al. 2014) and a ratiometric RGB camera system (Larsen et al. 2011) were used for O_2_ imaging of cross tissue sections of sheath-covered seagrass seeds. The O_2_ sensitive planar optodes were prepared via knife-coating a sensor cocktail onto a transparent polyethylene terephthalate (PET) foil. The O_2_ optode sensor cocktail consisted of 1.5 mg of platinum(II)-meso(2,3,4,5,6-pentafluoro)phenyl-porphyrin (PtTFPP; indicator dye), 1.5 mg of Macrolex® fluorescence yellow 10GN (MY; reference dye), 100 mg of polystyrene (PS), and 1 g of CHCL_3_. After dissolvement of all components in the CHCL_3_, the sensor solution was spread onto a dust-free PET foil using a film applicator (byk.com) yielding a ∼10 μm thick sensor film after evaporation of the solvent. To exclude background fluorescence and achieve the highest possible spatial resolution of the optical sensor, an optical isolation layer consisting of 1% w/w (10 mg) carbon black dispersed in 1 g 10% w/w solution of polyurethane hydrogel (D4) (EtOH: water, 9:1 w/w) was knife-coated on top of the optical sensor film yielding a final isolation layer thickness of ∼7.5 μm. The ratiometric RGB camera setup consisted of a SLR camera (EOS 1000D, Canon, Japan) equipped with a macro-objective lens (Macro 100 f2,8D, Tokina, Japan), and a 530 nm long-pass filter (uqgoptics.com). Excitation of the planar O_2_ optode was achieved via a 455 nm multichip LED (LedEngin Inc, RS Components Ltd, Corby, UK) combined with a bandpass filter. The excitation LED was powered by a USB-controlled LED driver unit (Triggerbox; imaging.fish-n-chips.de) interfaced with a PC running the custom software look@RGB (imaging.fish-n-chips.de) for image acquisition and control of the SLR camera and LED unit. The obtained RGB images were split into the red, green and blue channels, and analysed via the software ImageJ. For acquiring calibrated O_2_ concentration images, the red channel images (O_2_ sensitive emission of the indicator dye) and the respective green channel images (emission of the inert reference dye) were divided using the ImageJ plugin Ratio Plus (ratio = red/green); whereafter, the acquired ratio images were fitted to a previously obtained calibration curve (described in Figure S1) using the exponential decay function of the Curve Fitting tool in ImageJ.

### Data Analysis

The software program OriginPro 2017 (OriginLab Corporation, Northampton MA, US) was used for fitting and analysing the measured photosynthesis and respiration rates, and to obtain photosynthetic parameters as follows:

An exponential saturation model (Webb et al. 1974) was fitted for the gross photosynthesis (GP) rates as a function of photon scalar irradiance (E):

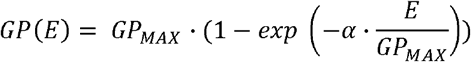

Similar exponential saturation model with an additional term to account for respiration (Spilling et al. 2010) was used for fitting net photosynthesis (NP) rates as a function of photon scalar irradiance (E):

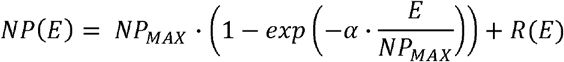

where *P_MAX_* is the calculated maximum photosynthesis rate at saturating photon irradiance, *α* is the light use efficiency related to the photosynthetic activity and efficiency, *R* is the respiration rate, and *E* is the respective photon scalar irradiance.

The compensation photon scalar irradiance (E_C_); i.e., the irradiance above which net photosynthetic production of O_2_ occurs, was determined from the achieved photosynthetic parameters as:

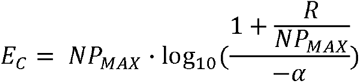

And finally, the photon scalar irradiance at the onset of photosynthesis saturation (E_K_) was calculated as:

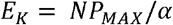

All oxygen production and consumption data were normalized to mg WW^-1^; to allow for utilizing the same seeds (i.e., as many as possible) for all measurements at the different development stages throughout the entire experiment, and thereby minimize biological differences between replicates.

## Results

### Photosynthesis and O_2_ microgradients in sheath-covered seeds

Variable chlorophyll fluorescence imaging revealed photosynthetic capacity in immature sheath-covered seagrass seeds (*Z. marina* L.), with an averaged maximum PSII photochemical quantum yield of 0.61 ± 0.04 (Fig. 1a; mean ± SD; n=3, biological replicates). Further development of the immature seed during opening of the seed sheath decreased the maximum PSII photochemical quantum yield to 0.50 ± 0.03 (Fig. 1b; mean ± SD; n=6, technical replicates), while seeds with detached sheaths displayed no photosynthetic capacity (Fig. 1c). Rapid light curves (RLC), i.e., measurements of the effective PSII photochemical quantum yield (YII) and relative photosynthetic electron transport rates (rETR) at increasing irradiance, further confirmed photosynthetic activity in *Z. marina* seed sheaths (Fig. 1d; n=3, biological replicates) following a typical saturation curve with increasing irradiance. Interestingly, mature *Z. marina* seeds without seed sheath also exhibited photosynthetic capacity, albeit with a markedly lower maximum PSII photochemical quantum yield of 0.28 ± 0.04 and a more patchy distribution of activity over the seed surface (Fig. 1c; mean ± SD; n=4, technical replicates).

**Figure 1.**
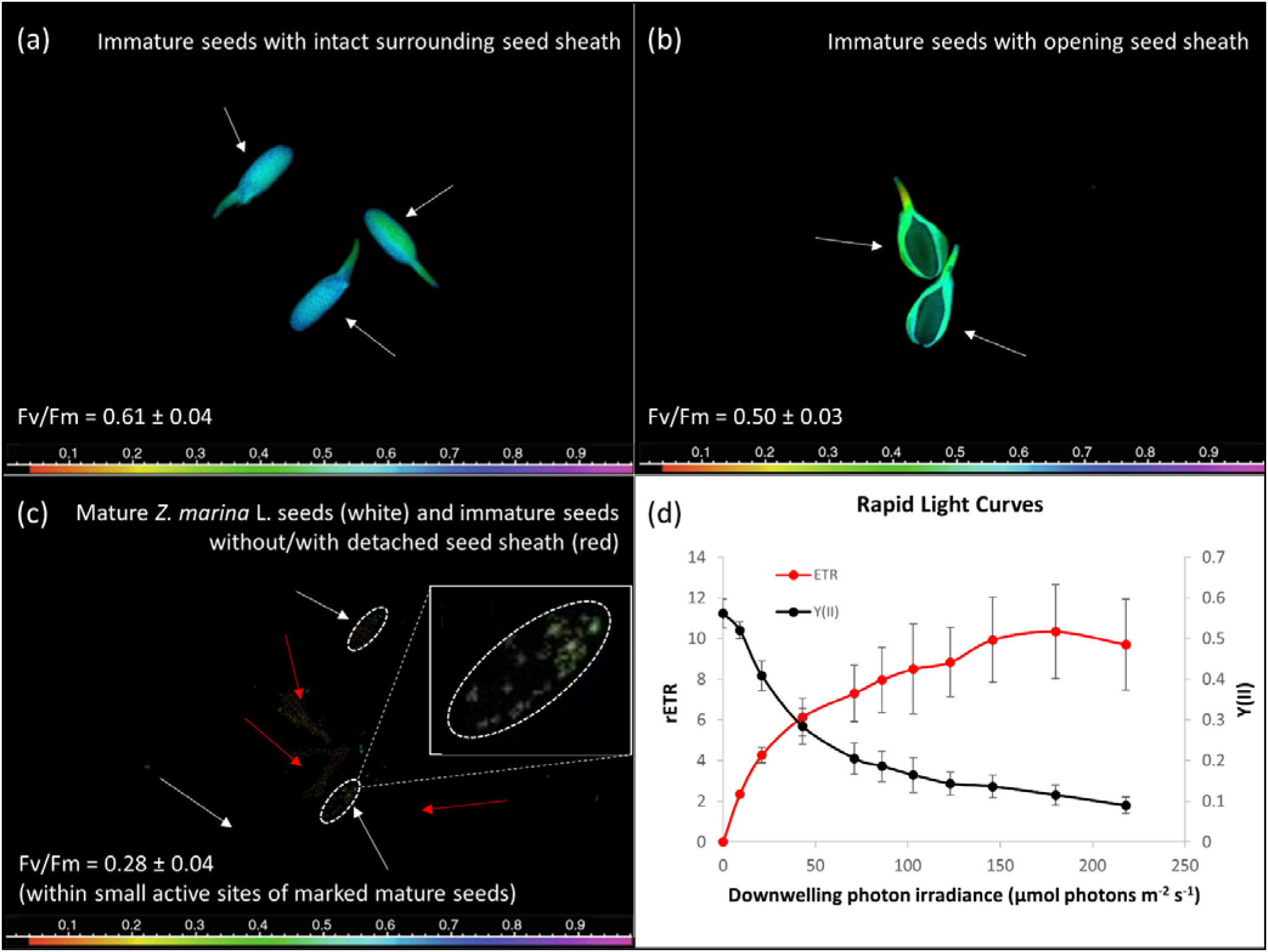
Photosynthetic capacity of seagrass seeds during early development stages. Variable chlorophyll fluorescence imaging of the maximum quantum yield of PSII (F_v_/F_m_; *i*.*e*., a measure of the PSII photochemical efficiency) of seagrass (*Z. marina* L.) seeds. (a) F_v_/F_m_ image of immature sheath-covered seeds of *Z. marina* (day 0). (b) F_v_/F_m_ image of immature *Z. marina* seeds with an opening seed sheath (day 3). (c) F_v_/F_m_ image of immature *Z. marina* seeds (*i*.*e*., white coloured seed coat) and mature *Z. marina* seeds (*i*.*e*., brown coloured seed coat) (day 6). Please note the small photosynthetic active regions of the mature seeds (marked within dashed white circles). (d) Rapid Light Curves (RLC) of immature, sheath-covered seeds of *Z. marina* (day 0), showing the relative electron transport rate (rETR) and the effective quantum yield of PSII [Y(II)]. n = 3, biological replicates.

Microscopic variable chlorophyll fluorescence imaging of cross tissue sections of sheath-covered immature *Z. marina* seeds showed that the photosynthetic capacity was solely restricted to the seed sheath (Fig. 2a). Photosynthesis activity of the seagrass seed sheath resulted in marked O_2_ production with increasing irradiance (Fig. 2b). At saturating irradiance, the maximal net and gross photosynthesis rates of seed sheaths were 6.08 ± 0.45 and 6.61 ± 0.44 nmol O_2_ mg WW^-1^ h^-1^, respectively, with light use efficiencies (α) under subsaturating irradiance of 0.10 ± 0.02 and 0.11 ± 0.03 (Table 1; mean ± SEM; n=6, biological replicates). The compensation photon irradiance (E_C_) and the onset of photosynthesis saturation (E_K_) of the seed sheaths were calculated to 30.1 and 62.9 μmol photons m^-2^ s^-1^, respectively (Table 1); indicating light requirements similar to seagrass leaves.

**Table 1.** Multiple photosynthetic parameters derived from the light response curves in figure 2, 4, 6 and 7. Data originate from seeds with intact seed sheath, seeds with epiphytes, surface-sterilized seeds, and young seedlings just after germination. Abbreviations refers to: P_n_ = net photosynthesis, P_G_ = gross photosynthesis, α = initial slope of the PI-curve in the light-limiting phase, P_max_ = maximum net photosynthesis rate, R = dark respiration rate, E_C_ = compensation photon irradiance, and E_k_ = onset of photosynthesis saturation.

|  | α | P<sub>max</sub><br>nmol O<sub>2</sub> mg WW<sup>-1</sup> h<sup>-1</sup> | R<br>nmol O<sub>2</sub> mg WW<sup>-1</sup> h<sup>-1</sup> | E<sub>C</sub><br>μmol photons m<sup>-2</sup> s<sup>-1</sup> | E<sub>k</sub><br>μmol photons m<sup>-2</sup> s<sup>-1</sup> |
| --- | --- | --- | --- | --- | --- |
| *Seeds with sheath* |  |  |  |  |  |
| P<sub>n</sub> | 0.10 ± 0.02 | 6.08 ± 0.45 | -4.06 ± 0.3 | 30.1 | 62.9 |
| P<sub>G</sub> | 0.11 ± 0.03 | 6.61 ± 0.44 |  |  |  |
| *Seeds with epiphytes* |  |  |  |  |  |
| P<sub>n</sub> | 0.13 ± 0.00 | 13.28 ± 0.25 | -2.82 ± 0.09 | 10.2 | 98.6 |
| P<sub>G</sub> | 0.17 ± 0.02 | 12.28 ± 0.82 |  |  |  |
| *Surface-sterilized seeds* |  |  |  |  |  |
|  |  |  | -2.8 ± 0.5 |  |  |
| *Seedling* |  |  |  |  |  |
| P<sub>n</sub> | 0.01 ± 0.00 | 3.23 ± 2.93 | -1.65 ± 0.17 | 125.9 | 405.0 |
| P<sub>G</sub> | 0.01 ± 0.00 | 1.98 ± 0.63 |  |  |  |

Values are means ± SEM of: (i) six seeds with intact seed sheath, (ii) three *Z. marina* L. seeds with epiphytes, (iii) four surface-sterilized seeds, and (iv) one seedling just after germination. Data originates from low O<sub>2</sub> conditions in the seawater to mimic the seeds/seedlings natural sediment environment (Fig. 4a,b, 6a,b and 7d); except for the surface-sterilized seeds dark respiration, as no O<sub>2</sub> production was found for surface-sterilized seeds during light illumination, and for seeds with sheath as they are normally growing in a naturally air saturated water-column environment. All data has been normalized to: mg WW<sup>-1</sup>.

**Figure 2.**
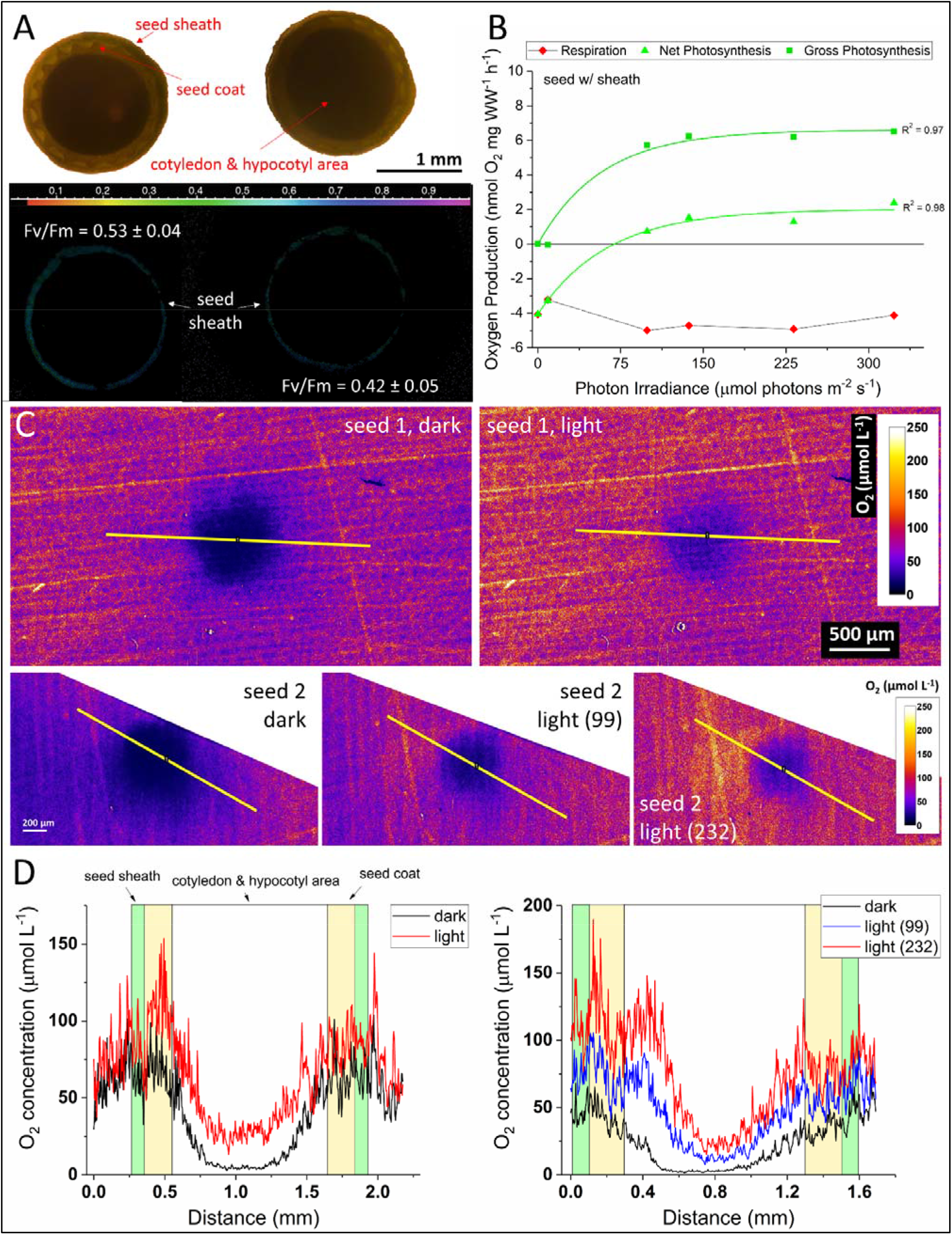
Photosynthetic capacity and architecture of sheath-covered seagrass seeds. (a) Photosynthetic capacity (F_v_/F_m_) over a cross section of a sheath-covered seed of *Z. marina* imaged with a variable chlorophyll fluorescence system mounted on an epifluorescence microscope. (b) Seed sheath O_2_ production and consumption rates as a function of increasing photon scalar irradiance (400-700 nm). The measured respiration, net- and gross photosynthesis rates are expressed as O_2_ evolution mg^-1^ biomass (WW). Rates of photosynthesis were fitted with an exponential saturation function (Webb et al. 1974), with an added respiration term (R) to account for respiration in the case of net photosynthesis rates (Spilling et al. 2010). R^2^ ≥ 0.97. Data originates from sheath-covered seeds of *Z. marina* (n=6). (c) Chemical imaging of sheath-covered seed with sheath cross section as shown in panel (a) via planar O_2_ optodes. The colour coded image shows the internal O_2_ concentration in sheath-covered seeds in light (99 and 232 μmol photons m^-2^ s^-1^) and darkness. (d) Extracted cross tissue line profiles (as shown in panel c), determining the internal O_2_ concentration gradient from the seed sheath, through the seed coat and into the cotyledon and hypocotyl tissue area of the *Z. marina* seeds. Black lines display the O_2_ concentration in darkness, blue line show the O_2_ concentration at a photon irradiance of 99 μmol photons m^-2^ s^-1^, and the red lines illustrates the O_2_ concentration at photon irradiances of 232 μmol photons m^-2^ s^-1^.

The dark respiration rate of sheath-covered seeds was -4.06 ± 0.3 nmol O_2_ mg WW^-1^ h^-1^ (Table 1; mean ± SEM; n=6, biological replicates), whereas post-illumination respiration, i.e., the respiration rate measured immediately after a light period under a defined irradiance, displayed a slight saturating increase with increasing irradiance (Fig. 2b). The photosynthetic activity resulted in increased internal O_2_ concentrations in the center of the immature seeds (i.e., the cotyledon and hypocotyl tissue area) enabling higher respiratory activity (Fig. 2c,d). Colour-coded O_2_ images of sheath-covered seed tissue cross-sections confirmed marked differences in the internal seed O_2_ concentration between measurements in light and darkness (Fig. 2c), where extracted line profiles across sheath-covered seeds showed a 5.9-fold increase in the internal O_2_ concentration from 4.6 to 27.3 μmol L^-1^ under a photon irradiance (400-700 nm) of 232 μmol photons m^-2^ s^-1^ in seed replicate 1 (Fig. 2d) and a 4.2- and 8.0-fold increase in the internal O_2_ concentration from 2.8 μmol L^-1^ in darkness to 12.1 and 22.7 μmol L^-1^ under photon irradiances of 99 and 232 μmol photons m^-2^ s^-1^, respectively, in seed replicate 2 (Fig. 2d). Hence, the photosynthetic O_2_ production of the seed sheath in the light is highly beneficial for the intra-seed O_2_ availability in developing immature *Z. marina* seeds (Fig. 2).

### Functional role of epiphytes for photosynthesis and respiration of mature seeds

Epiphyte cover on mature seagrass seed coats enabled active photosynthesis of the mature seed/epiphyte community (Figs. 1c and 3; Figs. S2 and S3). A biofilm of seed epiphytes covered most of the surface of the mature seed coat (Fig. 3a-d), which was dominated by diatoms but also harbored cyanobacteria, red and green algae (Fig. S4). The seed epiphyte layer had a maximum PSII photochemical quantum yield of up to 0.57 ± 0.01 (Fig. 3e; mean ± SD; n=6, technical replicates) and rapid light curves (RLC), i.e., measurements of the effective PSII photochemical quantum yield (YII) and relative photosynthetic electron transport rates (rETR) at increasing irradiance, showed photosynthesis saturation and inhibition with increasing irradiance (Fig. 3f).

**Figure 3.**
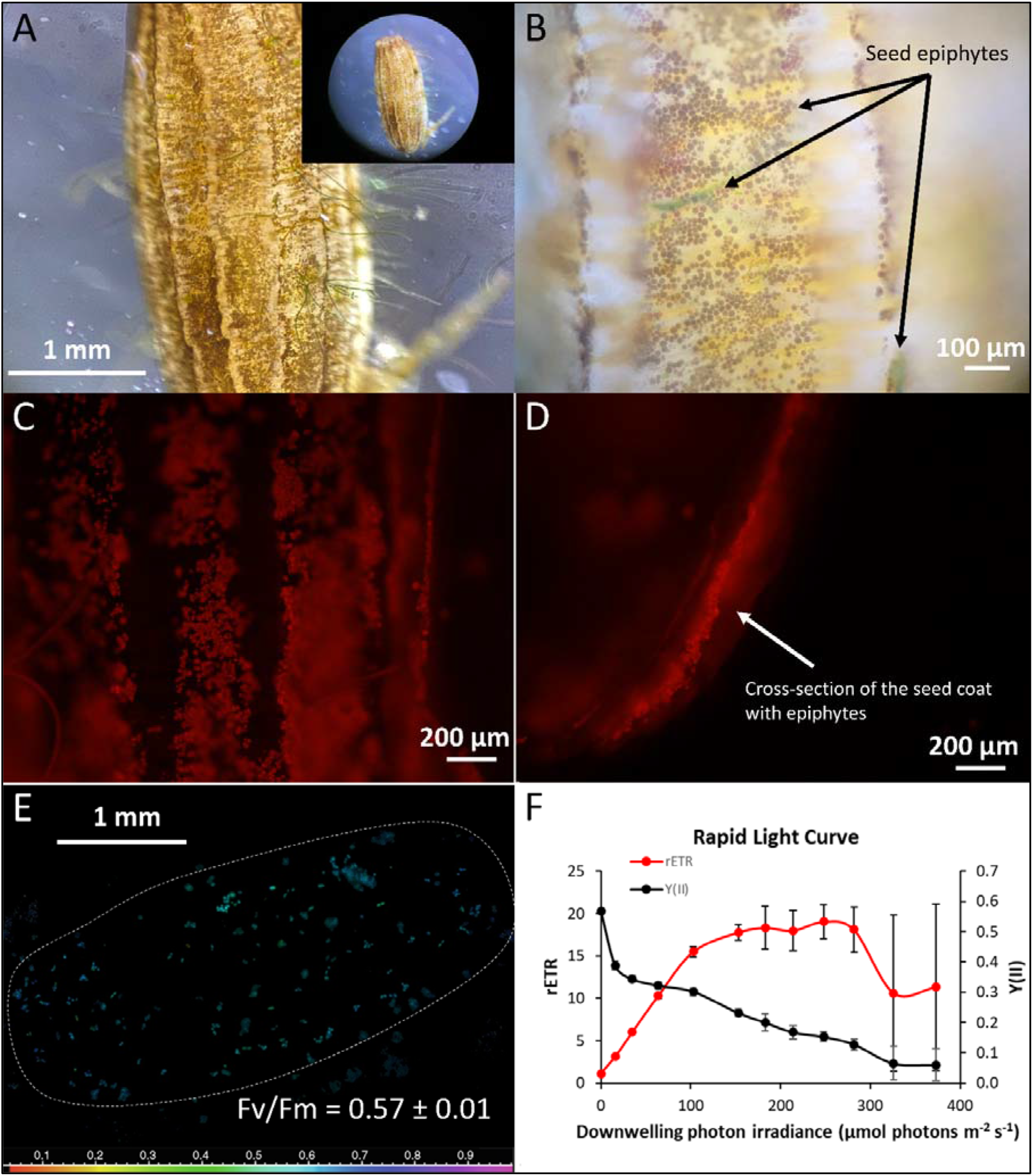
Photosynthetic capacity of mature seagrass seed with epiphytes. (a-b) Image of mature *Z. marina* seed showing the position of epiphytes on the seed coat. (c-d) Fluorescence microscope images of epiphyte chloroplasts, showing the seed coat surface (c) and a cross-section of the seed coat (d). (e) Microscopic variable chlorophyll fluorescence measurements of the PSII photochemical efficiency, i.e., the maximum quantum yield of PSII, F_v_/F_m_, of seed coat epiphytes (mainly consisting of diatoms, but also cyanobacteria, green and red algae, Fig. S4). The legend depicts the F_v_/F_m_ value. Further biological replication can be found in the supplementary Figs. S2 and S3. (f) Rapid light curves of the effective quantum yield of PSII [Y(II)] and the relative electron transport rate (rETR) of the seed epiphytes (n=3).

The photosynthetic activity of epiphytes on mature seeds led to marked O_2_ production and emission from the seed coat (Fig. 4). Initial gas exchange measurements during light/dark transitions revealed net photosynthesis rates of up to 15.8 ± 4.8 nmol O_2_ mg WW^-1^ h^-1^ and dark respiration rates of about -2.5 ± 0.6 nmol O_2_ mg WW^-1^ h^-1^ (Fig. 4a; mean ± SEM; n=6, technical replicates). Further detailed measurements and calculations of the photosynthetic O_2_ production of mature seed epiphytes under increasing irradiance determined maximal net and gross photosynthesis rates of 13.28 ± 0.25 and 12.28 ± 0.82 nmol O_2_ mg WW^-1^ h^-1^ at saturating irradiance with light use efficiencies under subsaturating irradiance of 0.13 ± 0.00 and 0.17 ± 0.02, respectively (Fig. 4b; Table 1; mean ± SEM; n=3, biological replicates). The compensation photon irradiance was calculated to 10.2 μmol photons m^-2^ s^-1^ and the onset of photosynthesis saturation first commenced at 98.6 μmol photons m^-2^ s^-1^ (Table 1). During darkness, the averaged dark respiration rate of the seeds with epiphytes was calculated to -2.82 ± 0.09 nmol O_2_ mg WW^-1^ h^-1^ (Table 1; mean ± SEM; n=3, biological replicates), whereas the post-illumination respiration increased to -4.0 nmol O_2_ mg WW^-1^ h^-1^ at a photon irradiance of 137 μmol photons m^-2^ s^-1^ followed by a drop in post-illumination respiration towards the highest measured irradiance of 232 μmol photons m^-2^ s^-1^ (Fig. 4b).

**Figure 4.**
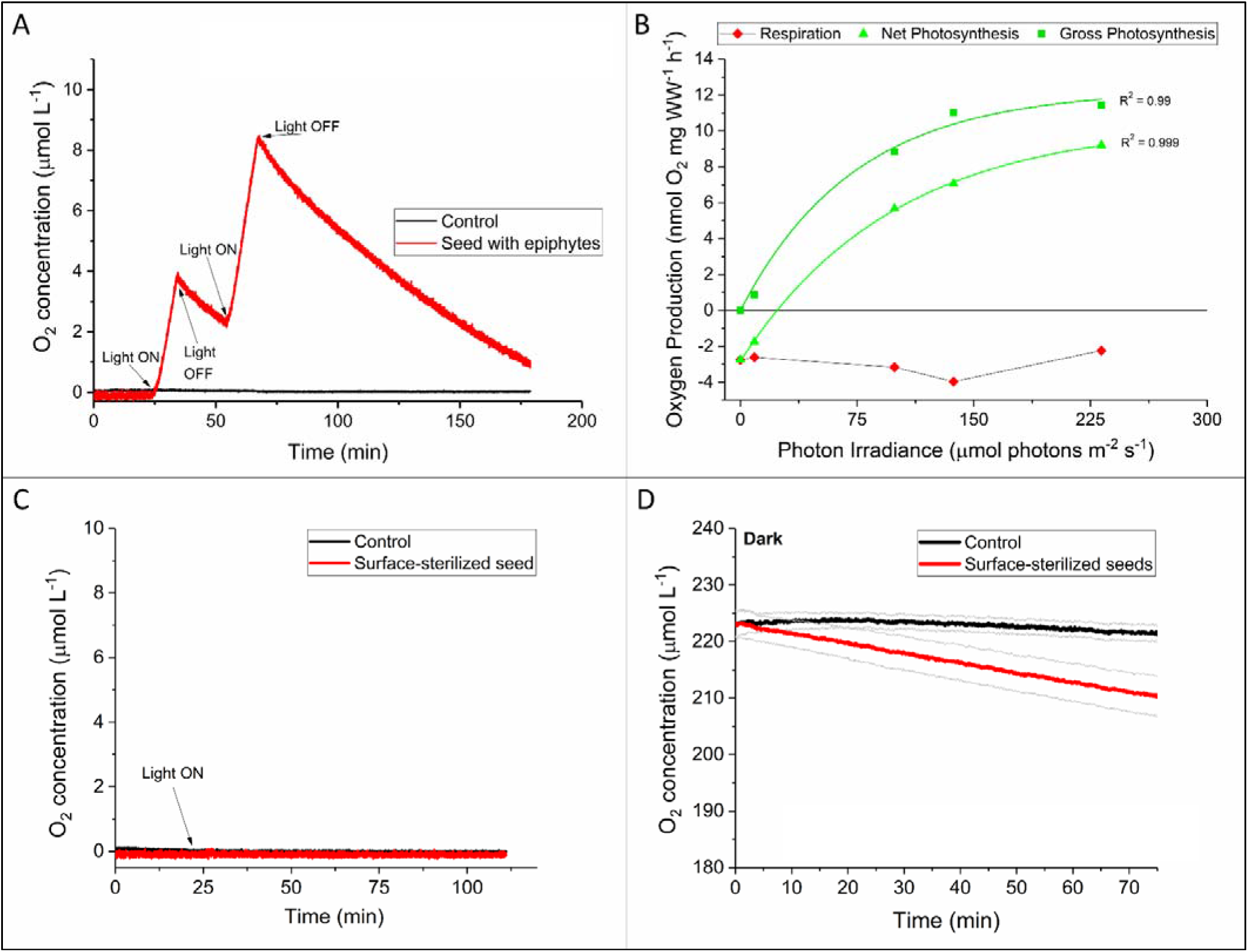
Photosynthetic O_2_ production and respiration rates of seagrass seeds with epiphytes and surface-sterilized seeds. (a) Net O_2_ production and consumption in seagrass seeds with epiphytes measured over time (photon irradiance of 232 μmol photons m^-2^ s^-1^). Averaged net photosynthesis and dark respiration rates were calculated from three *Z. marina* seeds with two technical replicates (n=6, technical replicates). (b) Seed with epiphytes photosynthesis and respiration as a function of photon scalar irradiance (400-700 nm). The measured respiration, net- and gross photosynthesis rates are expressed as oxygen evolution mg^-1^ biomass (WW). Measured photosynthesis values were fitted with an exponential saturation function (Webb et al. 1974), with an added respiration term (R) to account for respiration in the case of net photosynthetic rates (Spilling et al. 2010). R^2^ ≥ 0.99. Data originates from three *Z. marina* seeds with epiphytes (n=3). (c, d) Photosynthesis and dark respiration in surface-sterilized seeds. (c) Light response of surface-sterilized seed at a photon irradiance of 232 μmol photons m^-2^ s^-1^. Data shown for seed #1, while no O_2_ production in light was also found in seed #2 and #3 (n=3). (d) Dark respiration of surface-sterilized seeds (n=4). Black lines are control measurements (blank) and red lines show O_2_ measurements of surface-sterilized seeds. Gray lines show means ± SEM.

Surface-sterilized mature seeds showed no photosynthetic capacity and activity (Fig. 4c; n=3, biological replicates), but displayed similar dark respiration rates of -2.8 ± 0.5 nmol O_2_ mg WW^-1^ h^-1^ (Fig. 4d; mean ± SEM; n=4, biological replicates) as the mature seeds with epiphytes (Fig. 4a,b). The photosynthetic O_2_ production of mature *Z. marina* seeds is thus mainly driven by epiphytic diatoms, while the mature seeds themselves have relatively high respiratory needs.

### Photosynthetic capacity and O_2_ balance of early-stage seedlings of Z. marina

The photosynthetic capacity of early-stage *Z. marina* seedlings was investigated via variable chlorophyll fluorescence imaging and gas exchange measurements (Fig. 5). The basal hypocotyl (ROI 1), cotyledonary sheath (ROI 2), and cotyledonary blade (ROI 3) tissue area exhibited a maximum PSII photochemical quantum yield of 0.50, 0.44 and 0.30, respectively (Fig. 5a). The seedling tissue region with active root formation displayed no photosynthetic capacity (Fig 5a). Rapid light curves showing effective PSII photochemical quantum yields (YII) and relative photosynthetic electron transport rates (rETR) under increasing photon irradiance also confirmed photosynthetic activity in early-stage seedlings, where earlier photosynthesis inhibition with increasing irradiance was observed in the basal hypocotyl and cotyledonary sheath, as compared to the cotyledonary blade tissue area (Fig. 5b). Such photosynthetic capacity of the cotyledonary tissue was likely supported by the innermost growing first true leaf (Fig 5c). Microscopic variable chlorophyll fluorescence imaging of the cotyledonary blade apex showed that the photosynthetic capacity ceased at the tip of the cotyledonary blade (Fig. 5c). The main function of the cotyledonary blade apex may thus be sensing light and/or gravity. The variable chlorophyll fluorescence imaging also showed indications of minor and heterogenous epiphyte growth on the cotyledonary tissue surface (Fig. 5c). Measurements of photosynthetic O_2_ production and respiration of the early-stage seedling under increasing irradiance, revealed a relatively high respiratory demand and low photosynthetic activity of the seedling tissue with maximal net and gross photosynthesis rates of 3.23 ± 2.93 and 1.98 ± 0.63 nmol O_2_ mg WW^-1^ h^-1^ at high irradiance (Fig. 5d; Table 1; mean ± SE of the nonlinear curve fit; n=1, biological replicate). Light utilization efficiencies were also very low, i.e., 0.01 ± 0.00 for both net and gross photosynthesis rates (Fig. 1d; Table 1; mean ± SE of the nonlinear curve fit; n=1, biological replicate), leading to very high calculated compensation (E_C_) and onset of photosynthesis saturation (E_K_) irradiances of 125.9 and 405.0 μmol photons m^-2^ s^-1^, respectively (Table 1), as compared with seeds with sheath and epiphyte cover. The dark respiration rate of the early-stage seedling was -1.65 ± 0.17 nmol O_2_ mg WW^-1^ h^-1^ (Table 1; mean ± SE of the nonlinear curve fit; n=1, biological replicate). Early-stage seedlings are thus able to produce some O_2_ for their respiratory needs during growth, which likely is supported by photosynthesis of the innermost first true leaf; albeit, with much lower production rates as compared to the seed sheath photosynthetic capacity.

**Figure 5.**
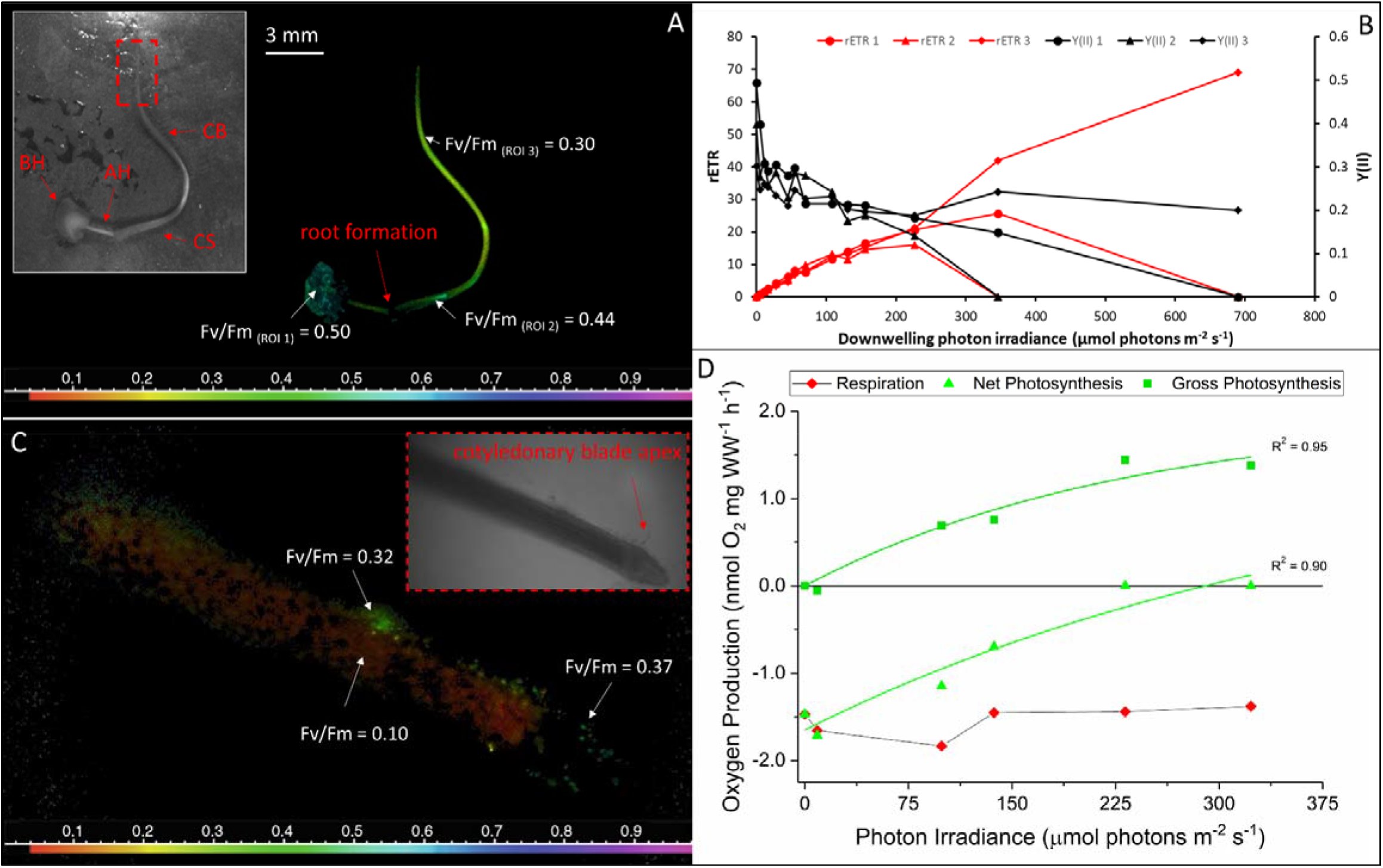
Photosynthetic capacity and oxygen production in seagrass seedling. (a) Maximum quantum yield of PSII (F_v_/F_m_) of young *Z. marina* L. seedling after germination. BH = basal hypocotyl (ROI 1), AH = axial hypocotyl, CS = cotyledonary sheath (ROI 2), and CB = cotyledonary blade (ROI 3). (b) Effective quantum yield of PSII [Y(II)] and relative electron transport rate (rETR) within the 3 selected regions of interest (ROIs 1-3) on the seedling as shown in panel A (n=3), determined at increasing irradiance via rapid light curves. (c) Microscopic variable chlorophyll imaging showing close-up image of the photosynthetic capacity (F_v_/F_m_) of the tip of the seedling cotyledonary blade with inner growing first true leaf. (A and C) Legend depicts the F_v_/F_m_ value. Grayscale pictures show the position of the seedling (*Z. marina* L.) and the cotyledonary blade apex. (d) Oxygen production and respiration rates of the seedling at increasing irradiance. The measured respiration, net- and gross photosynthesis rates are expressed as O_2_ evolution mg^-1^ biomass (WW). Photosynthesis values were fitted with an exponential saturation function (Webb et al. 1974), with an added respiration term (R) to account for respiration in the case of net photosynthetic rates (Spilling et al. 2010). R^2^ ≥ 0.9.

## Discussion

Our results provide first experimental evidence that seagrass (*Z. marina* L.) seed sheaths produce O_2_ in the light, leading to markedly increased internal O_2_ availability inside the cotyledon and hypocotyl tissue area of the seed (i.e., embryo), which ensures sufficient O_2_ support for respiration and thereby energy for biosynthetic activity in the developing seeds.

### Photosynthetic capacity and internal O_2_ gradients in developing seagrass seeds

Illumination of *Z. marina* seed sheaths led to strong gradients of seed photosynthesis and internal O_2_ concentrations in the immature seagrass seeds, from relatively high O_2_ production and concentration areas in the chlorophyll-containing and photosynthetic active surrounding seed sheath to hypoxic conditions in the inner embryo and endosperm tissue. After opening and detachment of the seed sheath, the photosynthetic capacity of the seed sheath ceased. Such seed sheath photosynthetic O_2_ production likely acts to relieve severe internal hypoxic conditions and stress in the developing seagrass seed, which may increase the supply of respiratory energy and thereby positively affect the seeds biosynthetic activity and thus carbohydrate reserves stored in the endosperm (Borisjuk and Rolletschek, 2009). Such O_2_ evolution capacity in sheath-covered seagrass seeds, is similar to observations in immature barley caryopsis seeds, which exhibit i) net O_2_ production at saturating light conditions (∼350 μmol photons m^-2^ s^-1^), ii) gross photosynthesis rates of 1.8 μmol O_2_ grain^-1^ h^-1^ and respiratory O_2_ demand of 0.4 μmol O_2_ grain^-1^ h^-1^, and iii) cessation of photosynthetic activity during maturation (Tschiersch et al. 2011). The seagrass seed coat and sheath thus serve several physiological functions that have evolved to promote the development of the seed. Here, seed sheath photosynthetic activity plays a vital role for endosperm storage activity, while the seagrass seed coat provides protection for the mature seed, which enables dormancy and seed dispersal and thus is of vital importance for successful establishment of seedlings in adjacent sediment areas.

### Impacts of epiphytic photosynthesis on the success of seed germination

Surface-sterilized, mature seagrass seeds did not exhibit photosynthetic capacity, however, showed sustained respiratory activity. Seed coat epiphytes (mainly diatoms) led to photosynthetic O_2_ production on the seed coat surface, which slightly enhanced post-illumination respiration rates of mature seagrass seeds, while it had no effects on the dark respiration rate. Whether such epiphyte-induced O_2_ production supports the mature seeds respiratory demand and thereby biosynthetic activity remains unknown and deserves further attention in future studies.

Epiphytic photosynthesis on the seed coat of mature seeds could also play a protective role via oxidation of phytotoxic compounds such as H_2_S during seed germination in the sediment surface layers. Such plant-derived oxic microhabitats in the sediment have been shown to be very important for seagrasses, as it functions as a chemical and biological defence mechanism in the seagrass rhizosphere against toxic H_2_S intrusion (Brodersen et al. 2015; 2018), and can also solubilise essential nutrients for growth (Brodersen et al. 2017). Nevertheless, the observed respiratory need of mature seagrass seeds may explain why seeds germinate earlier in anoxic sediment as compared to more oxygenated sediment conditions (e.g., Moore et al. 1993; Kawasaki, 1993, Orth et al. 2000), simply as a necessity for ensuring sufficient O_2_ support and respiratory energy for biosynthetic activity during germination and establishment of the seedling. Such demand for respiration energy in mature seed is supported by previous findings of improper development and reduced survival of *Z. marina* seeds exposed to anaerobic conditions (Hootsmans et al. 1987; Churchill, 1992).

### Comparison of metabolic activity in seeds and early-stage seagrass seedlings

Early-stage *Z. marina* seedlings showed photosynthesis capacity and activity of the hypocotyl and cotyledonary tissues, except at the site of adventitious root formation and at the apex of the cotyledonary blade. Generally, the cotyledonary tissue of monocotyledons (grass-like flowering plants such as seagrasses) are not considered photosynthetic, in contrast to dicot seedlings whose cotyledons display photosynthetic capacity (Ampofo et al. 1976; Harris et al. 1986; Brown & Huber, 1987; Zheng et al. 2011). The higher photosynthetic capacity in the cotyledonary sheath, as compared to the cotyledonary blade, could be derived from the innermost growing first true seagrass leaf, which elongates from the inside if the cotyledonary sheath (Xu et al. 2016). Hypocotyl photosynthesis has been shown in other plants, like in the green hypocotyl tissue of pine seedlings but is considered less important than cotyledon photosynthesis for seedling growth and development (Sasaki & Kozlowski, 1970).

The importance of cotyledon and hypocotyl photosynthesis on seedling development has largely been attributed to mobilization of stored reserves to support early seedling growth (Ampofo et al. 1976; Harris et al. 1986; Brown & Huber, 1987; Zheng et al. 2011). In soybeans, the cotyledons synthesize chlorophyll upon emergence from the soil, and cotyledon photosynthesis contributes carbon via assimilation to the developing seedling (Brown & Huber, 1987), albeit cotyledon photosynthesis rates are very low as compared with the true leaves (Harris et al. 1986). In castor seedlings, cotyledon photosynthesis provides carbohydrate and energy for the first true leaf to appear and maintains seedling growth until the first true leaf has expanded (Zhang et al. 2011). In castor seedlings, the cotyledon photosynthesis ceases during seedling development, however, cotyledon photosynthesis is sufficient to balance respiratory losses during early-stage seedling establishment (Zhang et al. 2011). Cotyledon photosynthates are essential for leaf production in *Acer*, where the photosynthates is first exported to the developing first leaf and subsequently to the hypocotyl and roots (Ampofo et al. 1976). Thus, similar functions of the determined cotyledon and hypocotyl photosynthesis in early-stage seagrass seedlings is likely. We found no photosynthetic capacity in the cotyledonary blade apex and we speculate that this region of the cotyledonary tissue mainly plays a role for light and gravitropic responses during early-stage seagrass seedling growth, as observed in *Arabidopsis* where photoreceptors regulate the development of hypocotyls and cotyledons as a response to light conditions (Sullivan & Deng, 2003) and generally in higher plant coleoptiles and root caps containing statocytes that are involved in graviperception (Raven & Edwards, 2001).

Early-stage *Z. marina* seedlings exhibited lower photosynthetic efficiency (α), maximum net photosynthesis rate (P_max_), and respiration rate (R), as compared with *Z. marina* seeds with sheath, which resulted in markedly higher photosynthesis saturation (E_k_) and compensation photon irradiance (E_c_) of the young seedling. The respiration rate and compensation photon irradiance of the immature sheath-covered seeds were markedly higher (i.e., 1.4- and 3-fold, respectively) indicating high metabolic activity in the developing seagrass seeds. Hence, the functioning of the seagrass seed sheath photosynthesis seems to be like in other angiosperms by enhancing the endosperm storage activity, while the development and growth of the young seagrass seedlings is supported by both cotyledon and hypocotyl photosynthesis, as well as, by stored energy reserves.

In summary, we have demonstrated that the sheath of seagrass (*Z. marina* L.) seeds has photosynthetic capacity, which results in increased O_2_ availability for the plant embryo in the light and can alleviate intra-seed hypoxic stress conditions and allow for increased biosynthetic activity. The photosynthetic architecture of the immature seagrass seeds drives strong gradients of photosynthesis and O_2_ concentrations across the seed tissues, from high O_2_ levels in the photosynthetic active tissue of the seed sheath to low O_2_ levels in the non-photosynthetic embryotic center of the seed. Such photosynthetic capacity of the developing seagrass seed ceases during maturation with the detachment of the seed sheath. Early-stage seedlings also posses’ photosynthetic capacity, which seemed supported by the development of the first green leaf inside the cotyledonary sheath, where the lack of photosynthetic activity in the cotyledonary blade apex suggest that this part of the cotyledon may rather function as a sensing organ for e.g., light and/or gravity. The resulting increased respiratory energy could enhance nutrient transport and endosperm storage capacity of the seagrass seed and seedling, and may be vital for the successful transition from seed to established seedling and thereby for the long-term performance of seagrass meadows.

## Acknowledgements

The research was funded by grants from the Carlsberg Foundation (CF16-0899; KEB); the Villum Foundation (00028156; KEB), and the Independent Research Fund Denmark (DFF-8022-00301B; MK).

## Author Contribution

KEB planned and designed the research with support from MK. KEB performed the experiments. KEB processed and analysed the data. KEB wrote the manuscript with editorial help from MK. Both authors have given approval to the final version of the manuscript.

## Competing Interest

The authors declare no conflict of interest.

## Data Availability

The data that support the findings of this study are available from the corresponding author upon reasonable request.

## Supporting Information

Additional supporting information may be found in the online version of this article.

**Fig. S1:** Planar optode calibration plot and fitting.

**Fig. S2:** Photosynthetic capacity of seeds with epiphytes #2 and 3.

**Fig. S3:** Variable chlorophyll fluorescence and RLC of seeds with epiphytes #4-6.

**Fig. S4:** RGB function fits as indicator of the seed epiphyte community composition (n=3).

## Notes

### Competing Interest Statement

The authors have declared no competing interest.

