## Supporting Information for "Photosynthetic capacity in seagrass seeds and early-stage seedlings of *Zostera marina* L"

**Figure list:**

**Fig. S1:** Planar optode calibration plot and fitting

**Fig. S2:** Photosynthetic capacity of seeds with epiphytes #2 and 3.

**Fig. S3:** Variable chlorophyll fluorescence and RLC of seeds with epiphytes #4-6.

**Fig. S4:** RGB function fits as indicator of epiphyte community composition (n=3).

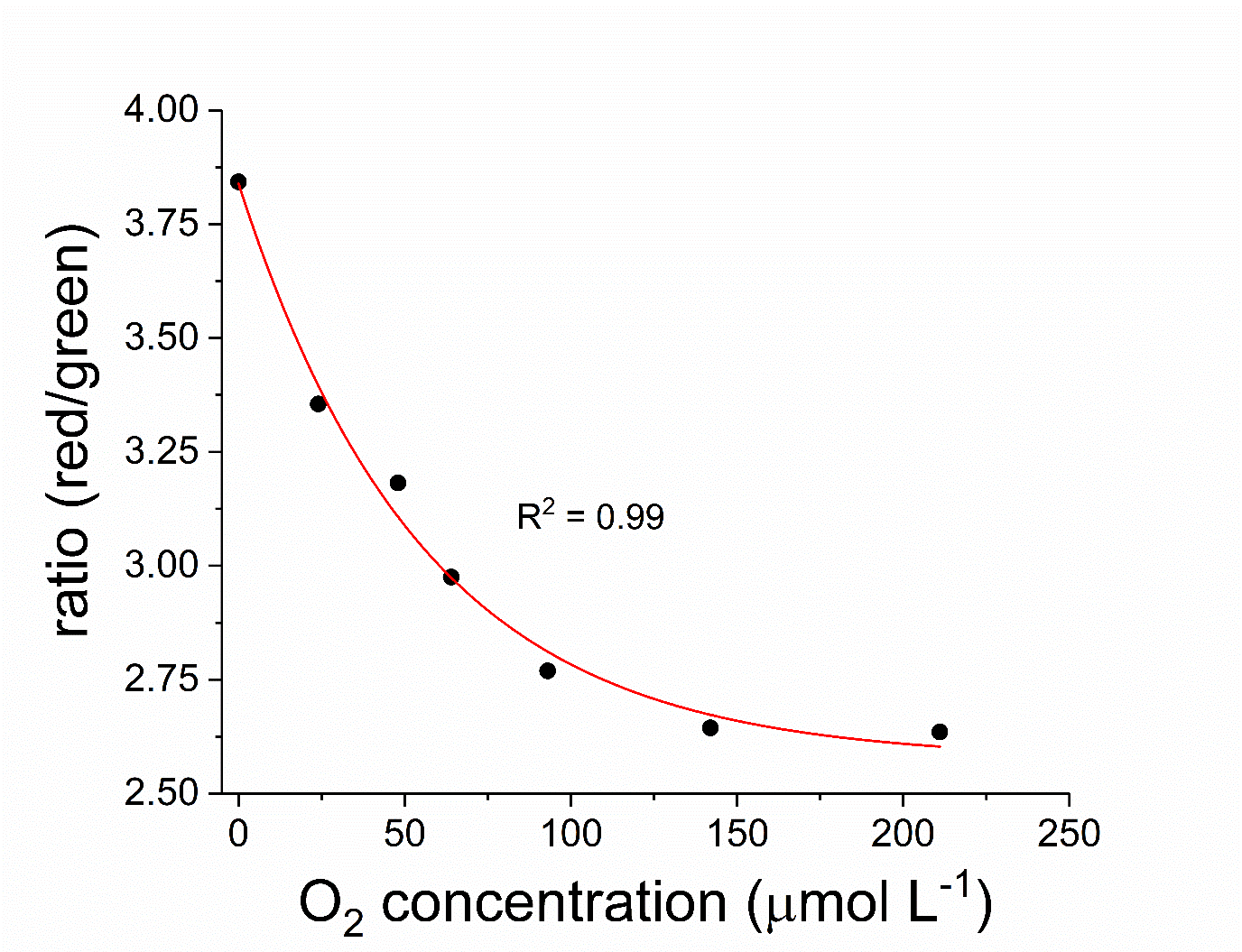

**Fig. S1. Planar optode calibration plot and fitting with exponential decay function.** The obtained exponential decay function (R^2^ = 0.99) was used to transform the acquired ratio images (i.e., red channel divided by green channel) to calibrated, colour coded O_2_ concentration images using the ImageJ software.

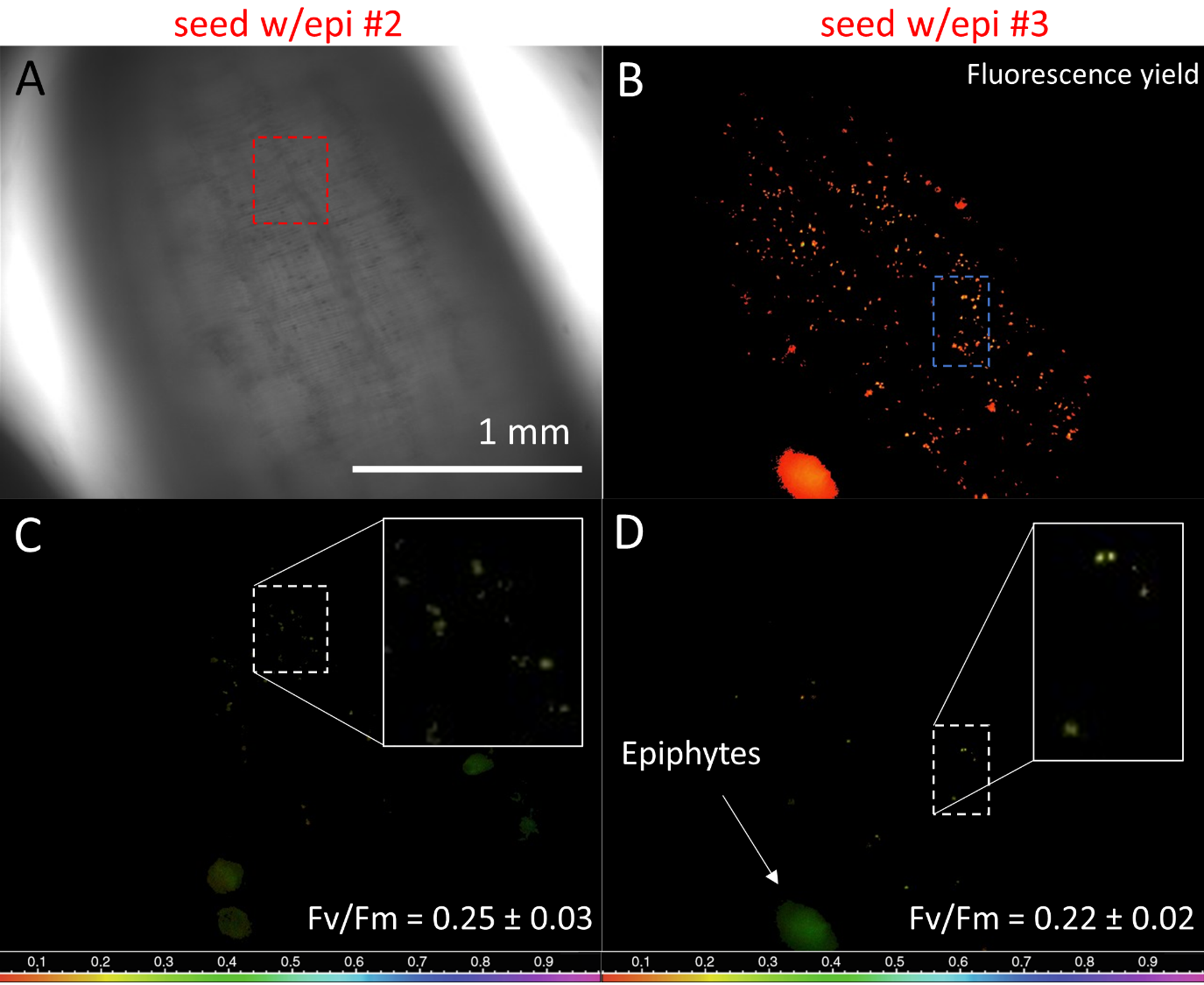

**Fig. S2. Photosynthetic capacity of seagrass seeds with epiphytes.** (A) Microscope image of seed coat surface. (B) Fluorescence yield measurement on the seed coat surface. (C) Variable chlorophyll fluorescence imaging of the maximum quantum yield of PSII (Fv/Fm). Panel C also shows a close-up image of the PSII maximum quantum yield on the seed coat for the region marked on panel A and C. (D) Variable chlorophyll fluorescence imaging of maximum quantum yields of PSII, with close up image of region displayed in panel B and D. Legend depicts the Fv/Fm value as color code (C-D).

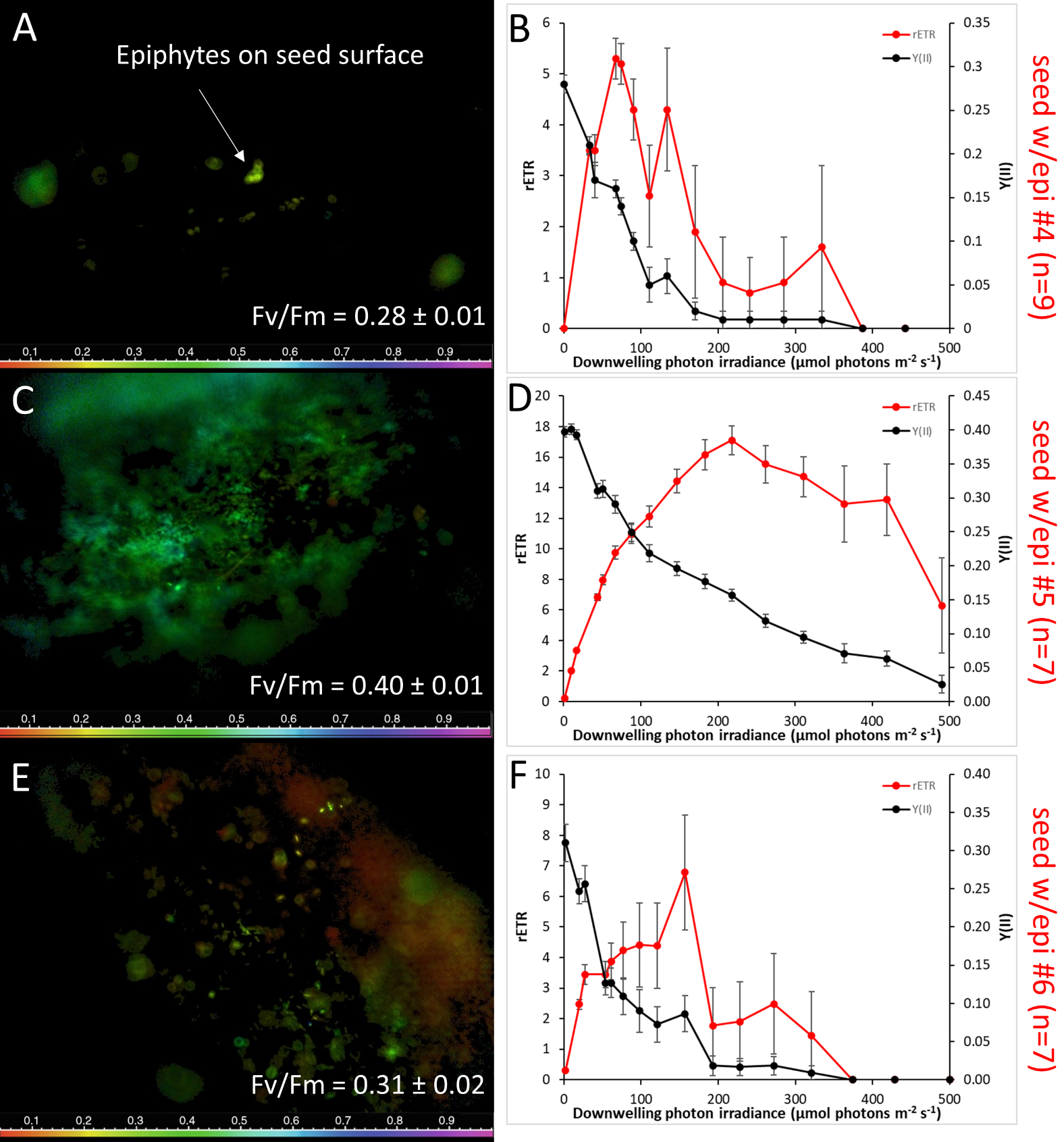

**Fig. S3. Variable chlorophyll fluorescence imaging and RLC of seeds with epiphytes.** The photosynthetic capacity expressed as maximum quantum yield of PSII (Fv/Fm) for seeds with epiphytes is shown in panel A, C, and E (*left panels*). Legend depicts the Fv/Fm value as color code. Rapid light curves (RLCs) of the relative electron transport rate (rETR) and the effective quantum yield of PSII (Y[II]) is shown in panel B, D and F (*right panels*). Values are means ± SEM (n = 7-9).

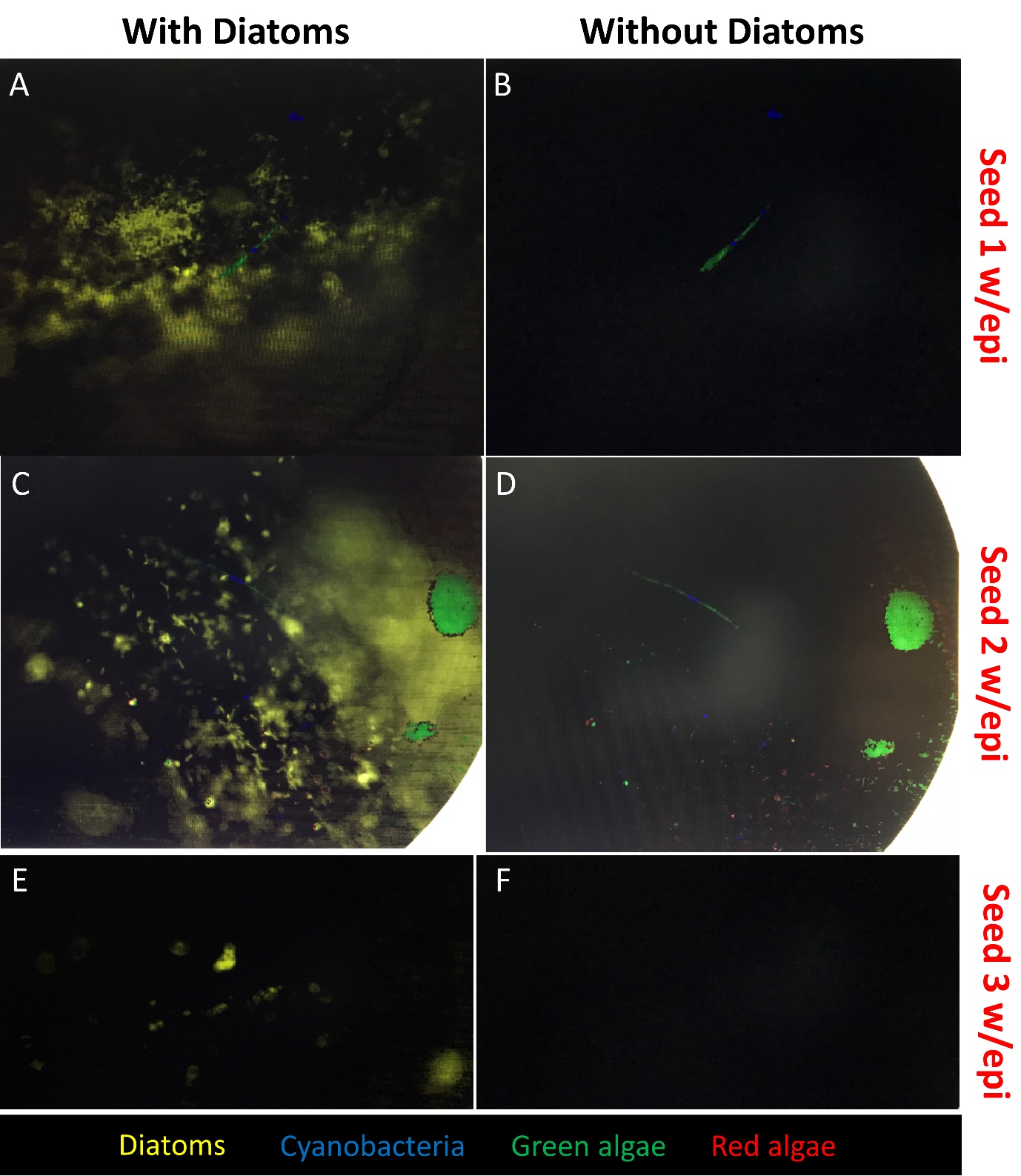

**Fig. S4. RGB function fits as indicator of epiphyte community composition on seagrass seeds.** Yellow color code indicates diatoms. Blue color code indicates cyanobacteria. Green color code indicates green algae. Red color code indicates red algae. Panels on the left are with diatoms, the dominating epiphyte species. Panels on the right are without diatoms. n=3.
